# ORA47 is a transcriptional regulator of a general stress response hub

**DOI:** 10.1101/2021.12.20.473540

**Authors:** Liping Zeng, Hao Chen, Yaqi Wang, Derrick Hicks, Haiyan Ke, Jose Pruneda-Paz, Katayoon Dehesh

## Abstract

Transcriptional regulators of general stress response (GSR) reprogram expression of selected genes to transduce informational signals into cellular events, ultimately manifested in plant’s ability to cope with environmental challenges. Identification of the core GSR regulatory proteins will uncover the principal modules and their mode of action in the establishment of adaptive responses. To define the GSR regulatory components, we employed a yeast-one-hybrid assay to identify the protein(s) that binds to the previously established functional GSR motif, coined Rapid Stress Response Element (RSRE). This led to the isolation of ORA47 (octadecanoid-responsive AP2/ERF-domain transcription factor 47), a Methyl jasmonate (MeJA) inducible protein. Subsequently, the ORA47 transcriptional activity was confirmed using RSRE-driven Luciferase (LUC) activity assay performed in the ORA47 loss- and gain-of-function lines introgressed into the 4xRSRE::Luc background. In addition, the prime contribution of CALMODULIN-BINDING TRANSCRIPTIONAL ACTIVATOR3 (CAMTA3) protein in induction of RSRE was reaffirmed by genetic studies. Moreover, exogenous application of MeJA led to enhanced levels of *ORA47* and *CAMTA3* transcripts, and the induction of RSRE::LUC activity. Metabolic analyses illustrated the reciprocal functional inputs of ORA47 and CAMTA3 in increasing JA levels. Lastly, transient assays identified JASMONATE ZIM-domain1 (JAZ1) as a repressor of RSRE::LUC activity.

Collectively, the report provides a fresh insight into the initial mechanistic features of transducing the informational signals into adaptive responses in part via the complex functional interplay between JA biosynthesis/signaling cascade and the transcriptional reprogramming necessary for potentiation of GSR, while offering a window into the role of intraorganellar communication in the establishment of adaptive responses.

**Significance:** The work unmasks the initial mechanistic features of adaptive responses that include tight cooperativity between JA biosynthesis and signaling cascade and the nuclear transcriptional machinery comprised of two activators (CAMTA3 and ORA47) and a suppressor JAZ1). The work further identifies CAMTA3 as a functional link between JA signaling and activation of a general stress transcriptional hub.

## Introduction

To survive organisms need to sense and rapidly respond to fluctuating environments, specifically by coordinating the necessary adaptive changes in metabolic pathways and physiological outputs. Stress-induced integrated transcriptional reprogramming of selected genes enables the rapid transduction of environmental signals into cellular responses, ultimately manifested in metabolic and physiological outputs crucial for coping with the imposed challenges.

The initial transcriptional reprograming known as the general stress response (GSR) is a recognized evolutionarily conserved stress response present across kingdoms (Marchler *et al*., 1993, HenggeAronis, 1996, Gasch *et al*., 2000, Chen *et al*., 2003, Kultz, 2005, Weber *et al*., 2005, Ma and Bohnert, 2007, Walley *et al*., 2007, Lopez-Maury *et al*., 2008, Walley and Dehesh, 2010, Benn *et al*., 2014, Benn *et al*., 2016). Previous studies of plants at 5 minutes post wounding led to the identification of an over-represented functional *cis*-element, dubbed the rapid stress response element (RSRE; CGCGTT), which is analogous to the yeast *stress response element* (*STRE*) (Kobayashi and McEntee, 1993, Marchler *et al*., 1993, Walley *et al*., 2007). Exploitation of transgenic Arabidopsis expressing RSRE::Luciferase (4xRSRE::LUC) confirmed the multi-stress responsive nature of RSRE induction and the suitability of the transgenic line for readout of stress induced rapid transcriptional responses (Walley *et al*., 2007, Bjornson *et al*., 2014, Benn *et al*., 2016, Benn and Dehesh, 2016). Additional molecular genetics and pharmacological approaches established the key role of the transcription factor, CALMODULIN-BINDING TRANSCRIPTION ACTIVATOR 3 (CAMTA3), in the induction of RSRE in a Ca^2+^ dependent manner (Bjornson *et al*., 2014, Benn *et al*., 2016).

Global transcriptomic profiling of Arabidopsis at 5 min post wounding identified several robustly induced genes encoding transcription factors, amongst them *ORA47* (octadecanoid-responsive AP2/ERF-domain transcription factor 47) (Walley *et al*., 2007). ORA47 belongs to a subfamily of the AP2/ERF domain transcription factors induced by methyl jasomonate (MeJA) (Nakano *et al*., 2006, Wang *et al*., 2008, Wasternack and Hause, 2013, Rehrig *et al*., 2014). ORA47 binds to the consensus motif (CCG(A/T)CC) in the promoters of various hormone biosynthesis genes, including the JA biosynthesis genes such as allene oxide synthase (*AOS*), allene oxide cyclases (*AOC1, AOC2, AOC3*), lipoxygenases (*LOX2 and 3*) and 12-oxo-phytodienoic acid (OPDA) reductase (*OPR3*) (Pauwels *et al*., 2008, Wasternack, 2015, Hickman *et al*., 2017).

The synthesis of jasmonate (JA) begins in chloroplast with LOXs, enzymes that convert ployunsaturated fatty acids to hydroperoxy derivatives. The hydroperoxy fatty acids are subsequently dehydrated into unstable allene oxides followed by their enzymatic conversion to 12-oxo-phytodienoic acid (OPDA) in the chloroplast (Creelman and Mullet, 1997). Next, OPDA is transported from chloroplasts to the peroxisomes where it is reduced by OPDA reductase (OPR3) and subsequently β-oxidized to yield JA, which is then conjugated to isoleucine to form JA-Ile in the cytosol (Howe, 2001, Wasternack and Hause, 2013). The jasmonate family (JAs) is comprised of JA, methyl jasmonate (MeJA), and OPDA, known signaling compounds with a pivotal role in a multitude of biological functions including metabolic processes, reproduction, responses to biotic and abiotic stresses, and transcriptional induction of the genes that regulate JA biosynthesis (McConn *et al*., 1997, Laudert and Weiler, 1998, Howe, 2001, Ishiguro *et al*., 2001).

Under standard conditions cells contain low JA levels, and transcription factors that induce JA-response genes are suppressed by the JASMONATE-ZIM DOMAIN (JAZ) family of transcriptional repressors (Chini *et al*., 2007, Thines *et al*., 2007, Yan *et al*., 2007). However, stress-induced biosynthesis of JA followed by JA-Ile production enable the interaction between JAZ proteins and the SCF^COI1^ ubiquitin ligase, leading to JAZ degradation via the 26S proteasome (Chini *et al*., 2007, Staswick, 2008, Chung *et al*., 2009). Destruction of JAZ transcriptional repressors lead to de-repression of multiple transcription factors and the activation of downstream JA response genes.

The search for GSR transcriptional regulators identified ORA47 as an activator, and JAZ1 as a suppresser of RSRE. The study also confirmed ORA47 functional involvement in increased 12-OPDA and enhanced *OPR3* expression accompanied by elevated JA levels. Collectively, the report unmasks the complexity of the intraorganellar communication network, projected through coordinated function of the phytohormone JA and the transcriptional regulators of a GSR element, ultimately enabling the ensued rapid transduction of environmental signals into accurate adaptive responses.

## Results

### *ORA47* binds to and induces RSRE activity

To isolate the RSRE-binding TFs, we performed a yeast-one-hybrid (Y1H) assay and identified several proteins amongst them ORA47 that binds to RSRE (Figure S1). To examine the transcriptional functionality of these RSRE-binding proteins in the induction of the *cis*-element, we employed TRANSPLANTA (TPT) collection of Arabidopsis lines that conditionally overexpress TFs under the control of a β-estradiol-inducible promoter (Coego *et al*., 2014). Next, we generated homozygous lines obtained from crosses between TPT plants expressing RSRE-binding proteins and the previously generated 4xRSRE::LUC line. Next we examined the LUC activity, before and after β-estradiol treatment of 4xRSRE::LUC lines in the wild type background (for simplicity, herein referred to as WT), and a number of TPT lines overexpressing RSRE-binding proteins found in the Y1H assay. However, based on functional analyses using RSRE-driven LUC activity assay before and after β-estradiol treatment, RSRE was exclusively induced in *TPT-ORA47* backgrounds (Figure S2a-b). The data clearly show a ~600-fold increase in LUC activity in the TPT-ORA47 lines post β-estradiol treatment, indicating that induction of ORA47 by β-estradiol activates RSRE-driven LUC activity.

To further examine RSRE-inducing activity of ORA47, we generated two independent *ORA47* CRISPR lines, designated *CRISPR-ora47-1* and −*2* in 4xRSRE::LUC line (Figure S3) using listed guide RNA sequences (Table S1). Next, we subjected these genotypes (WT, TPT-*ORA47*, *CRISPR-ora47-1* and −*2*) to mechanical wounding as an instantaneous and synchronous stimulus and examined their RSRE-driven LUC activity before and after wounding. Time course analyses show a rapid induction of the LUC activity, detectable at 5 min post wounding, peaking at 90 min in all lines albeit at different levels (Figure S4). The data further confirm that TPT-*ORA47* display constitutive bioluminescence signal under standard conditions, and a notably elevated wound-induced LUC activity as shown in a time course measurement assay (Figure S4). In addition, we also present representative images of seedlings at the peak LUC activity at 90 min post wounding, together with the corresponding measurements of LUC-activities (Figure 1a-b). Specifically, comparative activity analyses before and 90 min post wounding showed bioluminescence signals detected in unwounded TPT-*ORA47* at levels similar to those of the wounded WT plants. Moreover, wounding resulted in a ~30% higher bioluminescence signal intensity in TPT-*ORA47* compared to the WT plants. The basal bioluminescence measurements of the various genotypes detected similarly low signal intensity in *CRISPR-ora47-1* and −*2* and WT plants compared to that of TPT-*ORA47* lines (Figures 1a-b, S4). In fact, the basal bioluminescence signal in unwounded *TPT-ORA47* lines is similar to the WT levels albeit higher than those in CRISPR lines at 90 min post wounding. However, the easily detectable remanent of LUC signals in *CRISPR* lines, alluded to the contribution of transcription factor(s) other than ORA47 to the wound-induced RSRE-driven LUC activity (Figures 1a-b, S4).

**Figure 1.**
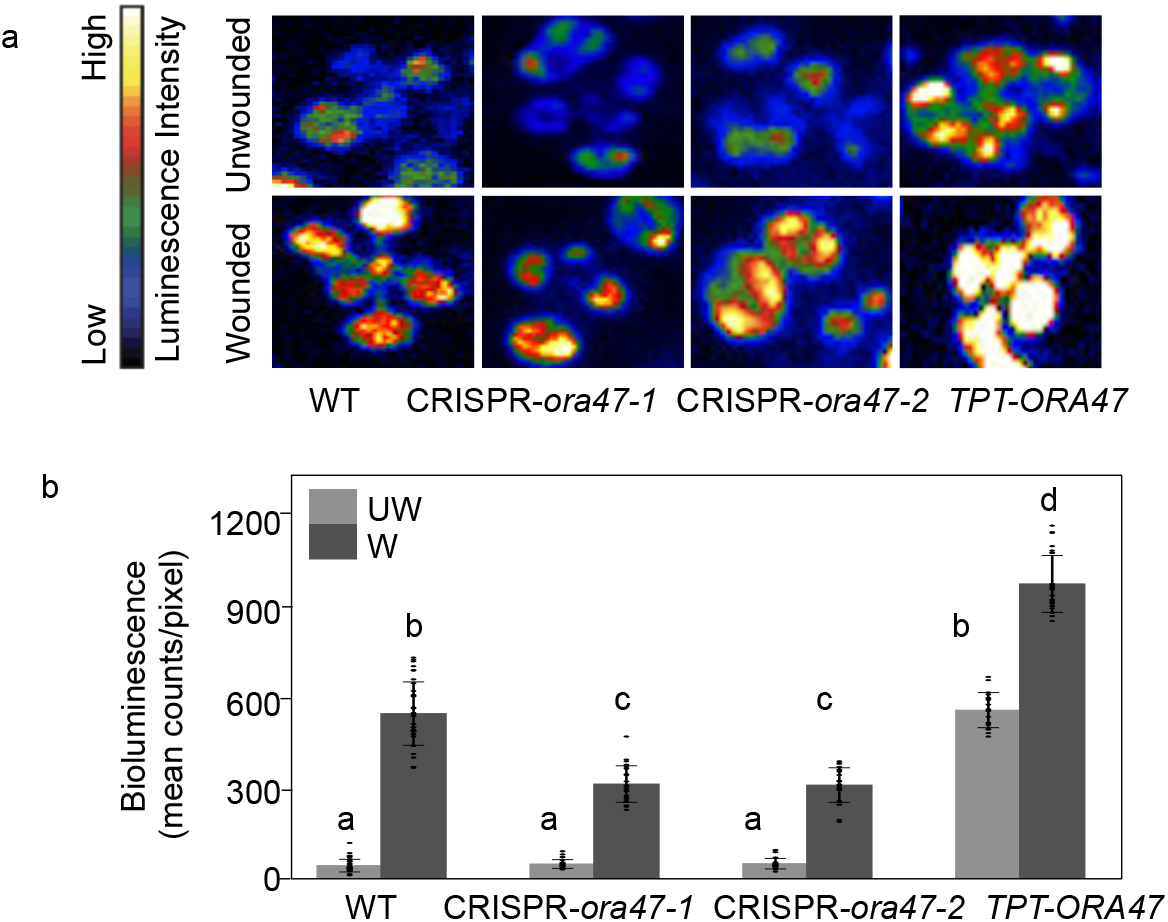
ORA47 induces RSRE::LUC activity. (a) Representative dark-field images of RSRE::LUC activity in unwounded (UW) and wounded (W, post 90 min) wild type (WT), *ORA47* loss of function mutants (*CRISPR-ora47-1* and *CRISPR-ora47-2*) and overexpression (TPT-*ORA47*) lines. The color-coded bar displays the intensity of LUC activity. (b) Quantitative measurements of LUC activity of plants shown in Panel A. Bars that do not share a letter represent statistically significant differences (p<0.05) by ANOVA test with Tukey’s honest significant difference (HSD) test. 36 plants per genotype per treatment were used as biological replicates. The error bar is the standard deviation of biological replicates.

### RSRE is induced by CAMTA3 and ORA47

The previously established function of CAMTA3 as an RSRE transcriptional activator (Bjornson *et al*., 2014, Benn *et al*., 2016), led us to specifically question the contribution of this protein to the RSRE::LUC activity in the *TPT-ORA47* genotype. Toward this goal, we crossed *TPT-ORA47* to *camta3* mutant and generated homozygous line (*TPT-ORA47/camta3*). Next, we compared the RSRE::LUC activity in unwounded and wounded (post 90 min) WT, *TPT-ORA47, camta3*, and TPT-*ORA47/camta3* lines (Figure 2a-b). The much reduced basal and wound-induced LUC activity levels in all *camta3* mutant backgrounds compared to their respective control lines confirmed the prime contribution of CAMTA3 to the induction of RSRE. Furthermore, the RSRE::LUC activity detected in wounded *camta3* mutant backgrounds, specifically in TPT-*ORA47/camta3*, supports the co-contribution of *ORA47* to the induction of RSRE.

**Figure 2.**
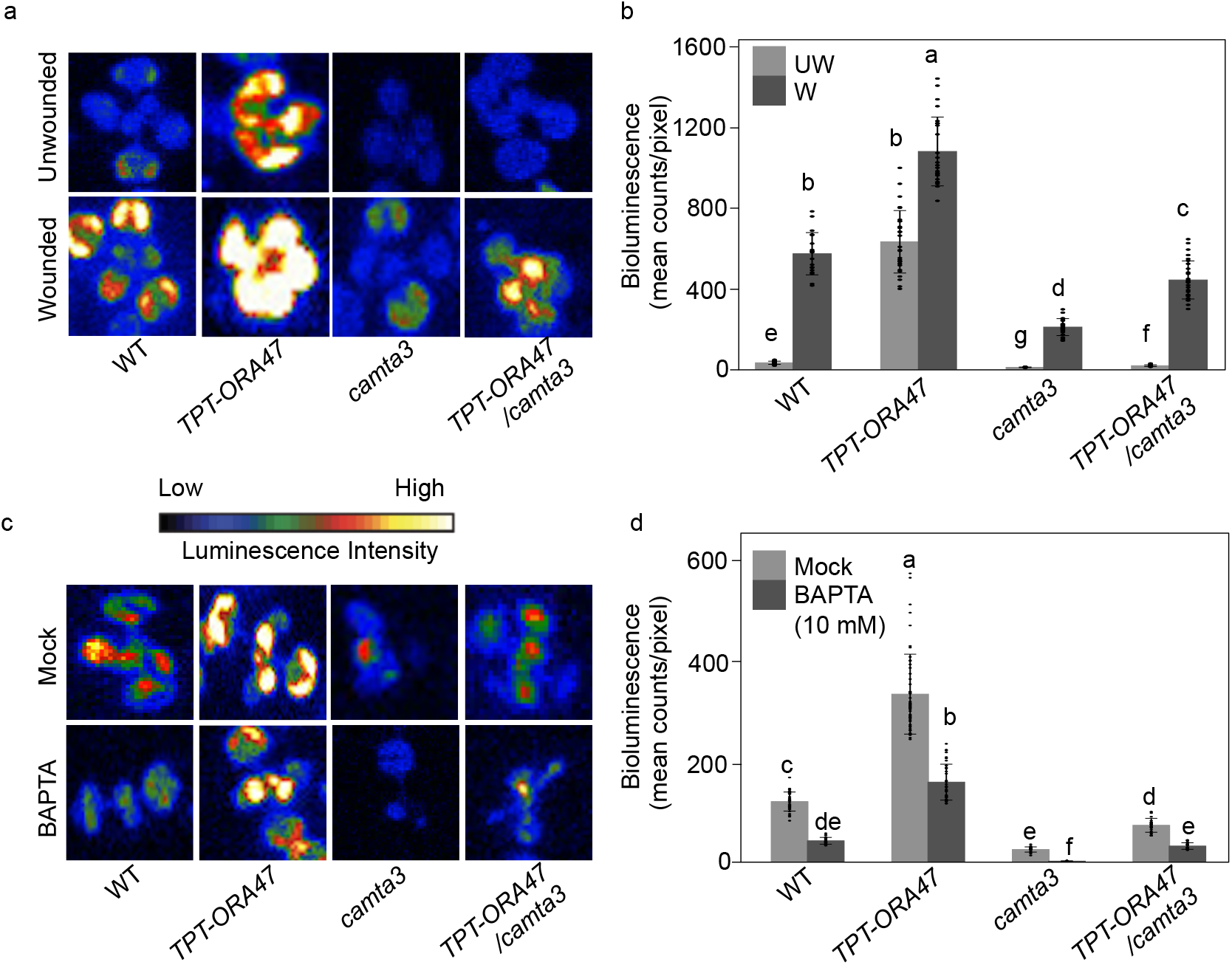
ORA47 induction of RSRE is Ca^2+^-dependent. (a) Representative dark-field images of RSRE::LUC activity in unwounded (UW) and wounded (W, post 90 min) wild type (WT), *TPT-ORA47, camta3*, and *TPT-ORA47/camta3* plants, and (b) their respective quantitative LUC activity measurements illustrate key function of CAMTA3 in RSRE induction. (c) Representative dark-field images of RSRE::LUC post 24 hr of mock- and BAPTA-treatment of aforementioned lines, and (d) their respective quantitative LUC activity measurements signify role of Ca^2+^ in ORA47-mediated induction of RSRE::LUC activity. The color-coded bar displays the intensity of LUC activity. Bars that do not share a letter represent statistically significant differences (p<0.05) by ANOVA test with Tukey’s honest significant difference (HSD) test. 50 plants per genotype per treatment were used as biological replicates for each treatment. The error bar is the standard deviation of biological replicates.

The reported key function of Ca^2+^ in CAMTA3 activation of RSRE (Benn *et al*., 2014, Benn *et al*., 2016), led us to question a potential role of this second messenger in ORA47 induction of this key GSR *cis*-element. To address this, we compared the RSRE::LUC activity in aforementioned genotypes treated with mock or with a selective Ca^2+^ chelator BAPTA (1,2-bis (o-aminophenoxy) ethane-N,N,N’,N’-tetraacetic acid) (Figure 2c-d). Highly reduced RSRE::LUC activity in all BAPTA-treated seedlings compared to those of the mock-treated lines illustrates that the ORA47 induction of RSRE is Ca^2+^-dependent. This data, together with the previous findings illustrating the Ca^2+^-dependent function of CAMTA3 in activation of RSRE (Benn *et al*., 2016, Bjornson *et al*., 2016) expands the functional repertoire of this second messenger in activation of GSR.

### MeJA induces RSRE activity

The reported MeJA-mediated induction of ORA47 (Wang *et al*., 2008, Rehrig *et al*., 2014, Chen *et al*., 2016) led us to explore the RSRE::LUC activity in mock- and MeJA-treated WT, TPT-*ORA47*, *camta3*, and *TPT-ORA47/camta3* seedlings. The analyses established MeJA-mediated induction of RSRE in all genotypes, most notably in TPT-*ORA47* followed by that of the WT, albeit with highly diminished bioluminance signal intensities in all the *camta3* mutant backgrounds compared to their respective controls, most notably in *TPT-ORA47/camta3* compared to the TPT-*ORA47* (Figure 3a-b). Moreover, the differential bioluminescence signal intensities in MeJA-/mock-treated versus those detected in wounded/unwounded TPT-*ORA47/camta3* seedlings (Figures 2a-b and 3a-b), allude to the potential function of CAMTA3 in the JA signaling cascade.

**Figure 3.**
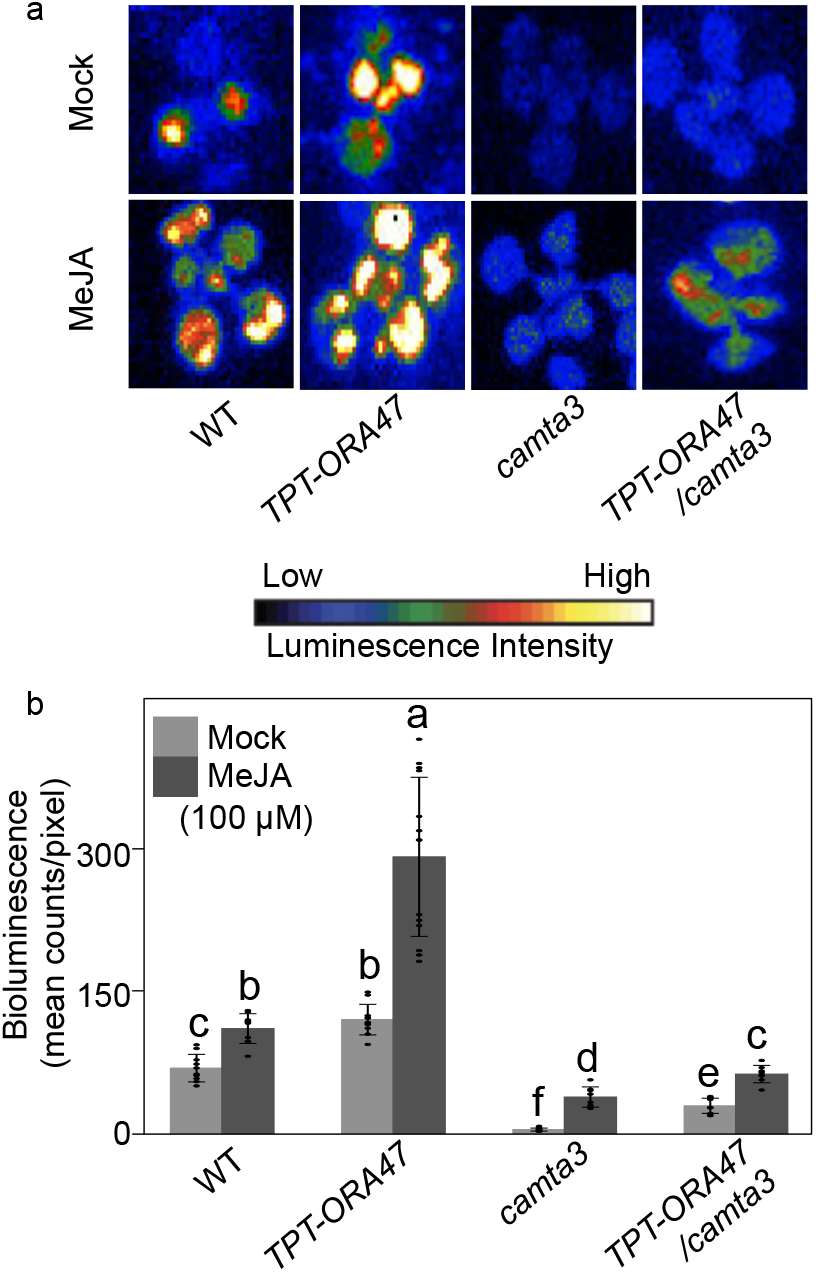
MeJA induces RSRE activity. (a) Representative dark-field images post 90 min Mock- and MeJA-treatment of WT, *TPT-ORA47, camta3*, and *TPT-ORA47/camta3* lines, and (b) their respective quantitative LUC activity measurements illustrate MeJA-mediated RSRE induction. The color-coded bar displays the intensity of LUC activity. Bars that do not share a letter represent statistically significant differences (p<0.05) by ANOVA test with Tukey’s honest significant difference (HSD) test. 36 plants per genotype per treatment were used as biological replicates. The error bar is the standard deviation of biological replicates.

### Positive feedback between JA, *ORA47* and *CAMTA3*

To assess potential contributions of CAMTA3 and ORA47 to the production of JA and its intermediate 12-OPDA, we analyzed the levels of these two metabolites in unwounded and wounded (90 min post wounding) WT, TPT-*ORA47*, *camta3*, and *TPT-ORA47/camta3* seedlings (Figure 4a-b). The data clearly show only slight wound-induced production of 12-OPDA exclusively in TPT-*ORA47* backgrounds (Figure 4a). In contrast however, wounding significantly enhanced JA production in all examined genotypes, particularly in TPT-*ORA47* lines, albeit with 40% reduction in *TPT-ORA47/camta3* line (Figure 4b).

**Figure 4.**
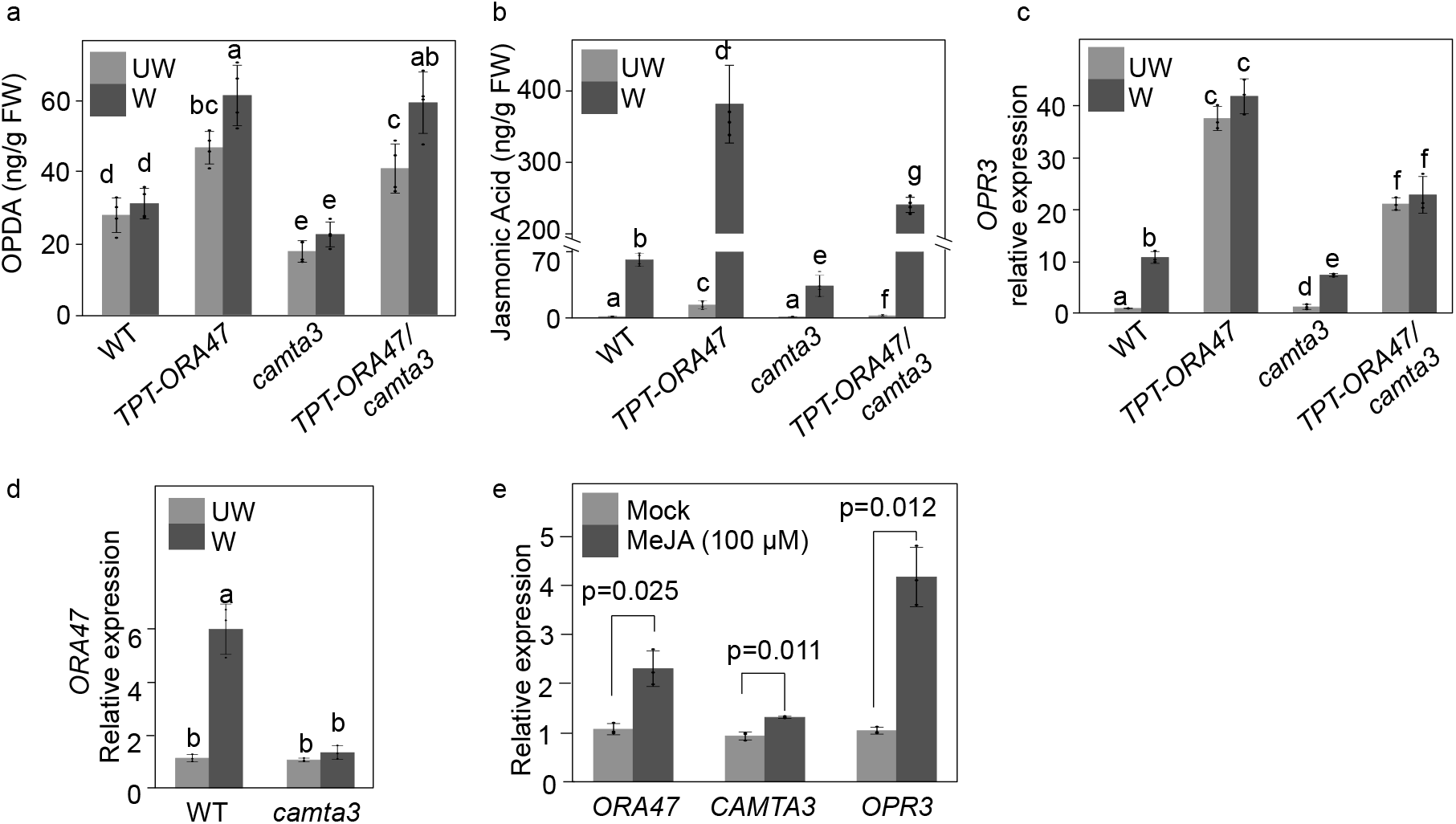
Positive feedback between JA and expression of *ORA47* and *CAMTA3*. (a) Analyses of 12-OPDA levels in 2-week-old unwounded (UW) and wounded (W, post 90 min) wild type (WT), *TPT-ORA47, camta3*, and *TPT-ORA47/camta3* plants. (b) Analyses of JA levels in aforementioned UW and W genotypes illustrate differential contribution of ORA47 and CAMTA3 to JA production. (c) Relative expression level analyses of *OPR3* in aforementioned genotypes display induction of the gene by ORA47 and CAMTA3. (d) Expression analyses of *ORA47* in UW and W wild type (WT) and *camta3* mutant lines illustrate *CAMTA3*-dependent expression of the gene in wounded plants. (e) Relative expression level analyses of *ORA47*, *CAMTA3*, and *OPR3* in mock- and MeJA-treated plants (post 90 min) illustrate MeJA induction of the genes. Bars that do not share a letter represent statistically significant differences (p<0.05) by ANOVA test with Tukey’s honest significant difference (HSD) test. The P value on top of the histograms were obtained by Student’s t-test. The error bar is the standard deviation of four to five biological replicates per genotype per treatment in (a) and (b), and three biological replicates in (c), (d), and (e).

The significantly higher wounding-induced JA levels in TPT-*ORA47* backgrounds relative to that of the WT is in concordance with the higher content of the intermediate, 12-OPDA, and by extension enhanced flux to JA production, potentially supported by elevated OPR3 for reduction of 12-OPDA in peroxisomes. To explore this possibility, we examined *OPR3* relative transcript levels in the aforementioned genotypes (Figure 4c). The data show that wounding results in increased *OPR3* transcript levels in WT and *camta3* mutants, but not in the TPT-*ORA47* backgrounds. The lower levels of *OPR3* transcripts in wounded *camta3* compared to the WT, suggest the involvement of CAMTA3 in transcriptional regulation of this gene. Indeed, the lower *OPR3* expression levels in *TPT-ORA47/camta3* compared to those of the TPT-*ORA47* line supports the transcriptional role of *CAMTA3* in induction of *OPR3*. However, the notably higher *OPR3* transcript levels in *TPT-ORA47* backgrounds versus the levels in WT and *camta3* lines also support the function of *ORA47* as an inducer of *OPR3* transcript levels. Furthermore, the wounding-independent increase in *OPR3* expression levels in *TPT-ORA47* backgrounds suggest that overexpression of *ORA47* is a substitution for an otherwise other wound-inducible regulators.

To examine the wounding-mediated transcriptional regulation of *ORA47* and the input of the *CAMTA3* in the process, we analyzed relative expression levels of *ORA47* in unwounded and wounded (90 min post wounding) WT and *camta3* mutant plants (Figure 4d). The data shows similar basal *ORA47* transcript levels in both genotypes, and further demonstrate exclusive wound-induced expression of the gene in the WT but not in *camta3* mutant. This alludes to functional input of *CAMTA3* in transcriptional regulation of *ORA47* in response to wounding.

Next, we examined possible JA-mediated transcriptional regulation of *ORA47, CAMTA3* and *OPR3* in mock- and MeJA-treated WT plants (Figure 4e). The data show basal expression levels of all genes in mock-treated plants compared to their higher relative expression in response to MeJA application, albeit at different degrees.

In summary, the above findings illustrate the intricacy of CAMTA3 and ORA47-mediated regulatory network that induce JA-biosynthesis pathway genes, and further show the reciprocal positive feedback of JA in induction of *CAMAT3* and *ORA47* transcript levels. Moreover, the data identifies CAMTA3 as a direct or an indirect positive regulator of *ORA47* expression.

### JAZ1 suppresses RSRE

To explore the underlying mechanism of MeJA-mediated induction of RSRE (Figure 3), we exploited tobacco transient expression assay to examine potential functional input of one of the JAZ transcriptional repressors on the RSRE-driven LUC activity. We specifically tested LUC activity in unwounded and wounded tobacco leaves transiently expressing JAZ1 alone and together with ORA47 and CAMTA3, individually (Figure 5a-d). The dark field images and their respective bioluminescence signal intensity measurements illustrate reduced basal and wound-inducible LUC activity in the presence JAZ1 when co-expressed with ORA47 (Figure 5a-b), and with CAMTA3 (Figure 5c-d). Accordingly, the data identifies JAZ1 as a suppressor of RSRE-activity and by extension a repressor of the ensued adaptive responses.

**Figure 5.**
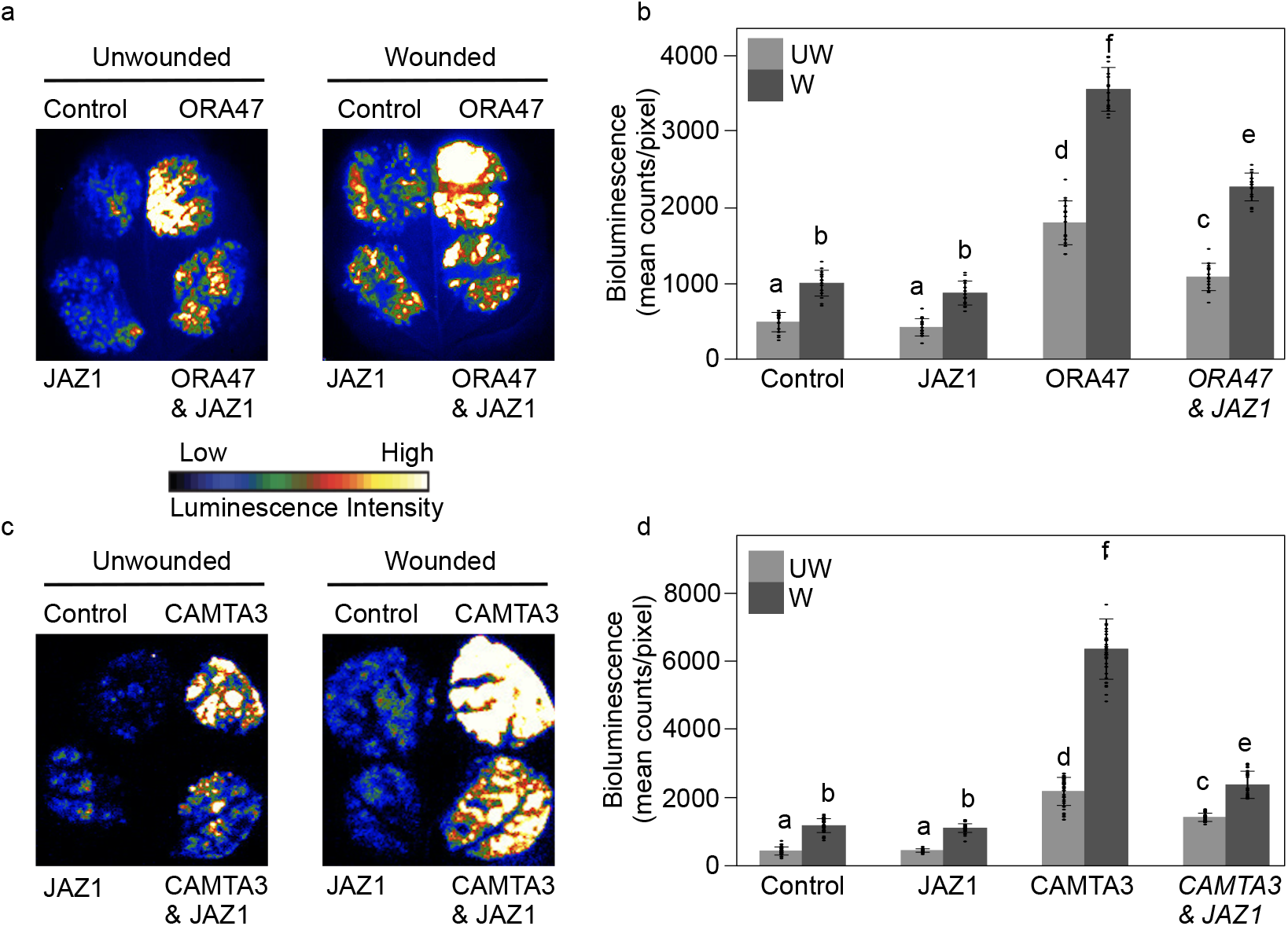
JAZ1 is a suppressor of RSRE. (a) Representative dark-field images of transiently expressed 35S-empty vector & RSRE::LUC (Control), 35S::JAZ1 & RSRE::LUC (JAZ1), 35S::ORA47 & RSRE:LUC (ORA47), 35S::ORA47 & 35S::JAZ1 & RSRE:LUC (ORA47 & JAZ1) in unwounded and wounded (post 90 min) tobacco leaves show JAZ1 suppression of ORA47-induced RSRE::LUC activity, and (b) the corresponding quantitative LUC activity measurements. (c) Representative dark-field images of transiently expressed 35S-empty vector & RSRE::LUC (Control), 35S::JAZ1 & RSRE:LUC (JAZ1), 35S::CAMTA3 & RSRE:LUC (CAMTA3), 35S::CAMTA3 & 35S::JAZ1 & RSRE:LUC (CAMTA3 & JAZ1) in unwounded and wounded (post 90 min) tobacco leaves illustrate JAZ1 suppression of CAMTA3-induced RSRE:LUC activity, and (d) the corresponding quantitative LUC activity measurements. Bars that do not share a letter represent statistically significant differences (p<0.05) by ANOVA test with Tukey’s honest significant difference (HSD) test. The error bar is the standard deviation of biological replicates. 20 leaves were used as biological replicates. The error bar is the standard deviation of biological replicates.

## Discussion

Stress-induced reprogramming of the GSR genes results in reconfiguration of selected stress signaling networks and the consequential biochemical and physiological output deemed for coping with environmental challenges. Defining the regulatory components of GSR provides a platform for delineating the underlying machinery involved in rapid and transient induction of early stress responses, thereby unmasking the initial mechanistic features of adaptive responses. Here, we utilized a rapidly and transiently activated multi-stress response *cis*-element, RSRE, present in ~30% of stress response genes (Yang and Poovaiah, 2002, Benn *et al*., 2016, Yuan *et al*., 2018), and identified ORA47 and CAMTA3 as two transcription factors that differentially induce this GSR transcriptional hub. Specifically, we genetically reaffirmed CAMTA3 as the prime inducer of RSRE, as previously shown (Doherty *et al*., 2009, Kim *et al*., 2013, Benn *et al*., 2016, Bjornson *et al*., 2016). This finding however is despite our inability to directly detect the physical binding of CAMTA3 to RSRE in the Y1H assay. The lack of a direct binding could be potentially caused by the absence of Ca^2+^/calmodulin, the suggested allosteric modulators of CAMTA3 (Benn *et al*., 2016), in the assay.

However, the Y1H assay established physical binding of ORA47 to RSRE. Binding of ORA47 to the previously identified consensus motif (CCG(A/T)CC) (Hickman *et al*., 2017), as well as its binding to the RSRE motif (CGCGTT) (Walley *et al*., 2007) illustrate the ability of this protein to bind to GC rich motifs albeit with some degree of promiscuity, ultimately resulting in induction of the JA biosynthesis genes and potentiation of initial stress responses by activation of selected GSR genes. Indeed, genetic analyses support the role of ORA47 in activation of RSRE, a key GSR transcriptional hub. In addition, ORA47 induction of RSRE, similarly to CAMTA3, is Ca^2+^ dependent. This finding expands the signaling role(s) of this second messenger in induction of a GSR transcriptional hub, albeit by yet an unknown mechanism.

In addition, the MeJA-mediated induction of RSRE activity tightly links the two rapidly stress response pathways whereby the stress induced production of JA and its bioactive-derivatives (<5 min) (Koo *et al*., 2009) promptly promote activation of a GSR transcriptional hub. Moreover, the dependency of this induction on CAMTA3 establishes the functional link between JA downstream signaling and activation of RSRE. One possible function of CAMTA3 is to induce JA production, and the consequential degradation of JAZ and the release of JAZ-mediated suppression of transcription factors, among them MYC2 and closely related bHLH TFs MYC3 and MYC4 that activate a large group of JA-responsive genes (Dombrecht *et al*., 2007, Fernandez-Calvo *et al*., 2011, Chen *et al*., 2016, Hickman *et al*., 2017, Van Moerkercke *et al*., 2019). One of the MYC2 targets is ORA47 that activates not only JA biosynthesis (Hickman *et al*., 2017), but also a GSR transcriptional hub. Indeed identification of JAZ1 as a suppressor of RSRE unmasks the underlying mechanism of JA action in activation of a GSR transcriptional hub through 26S proteasome-mediated degradation of JAZ (Chini *et al*., 2007, Staswick, 2008, Chung *et al*., 2009).

In summary, the schematic model (Figure 6) is a simplified depiction of a coordinated communication network between chloroplast/peroxisome/cytosol/nucleus, potentiating adaptive responses that begin with chloroplast-mediated increase in the available Ca^2+^ pool for activation of CAMTA3, followed by CAMTA3-mediated induction of *OPR3* and *ORA47* transcript levels and the consequential JA production. Next, JA production leads to degradation of JAZ1, an RSRE repressor, and reciprocal induction of *CAMTA3* and *ORA47*, the inducers of RSRE. Collectively, the data provide a window into the complexity of intraorganellar communication network facilitating the coordinated function of the phytohormone JA and the transcriptional regulators of a GSR element, ultimately enabling rapid transduction of environmental signals into accurate adaptive responses.

**Figure 6.**
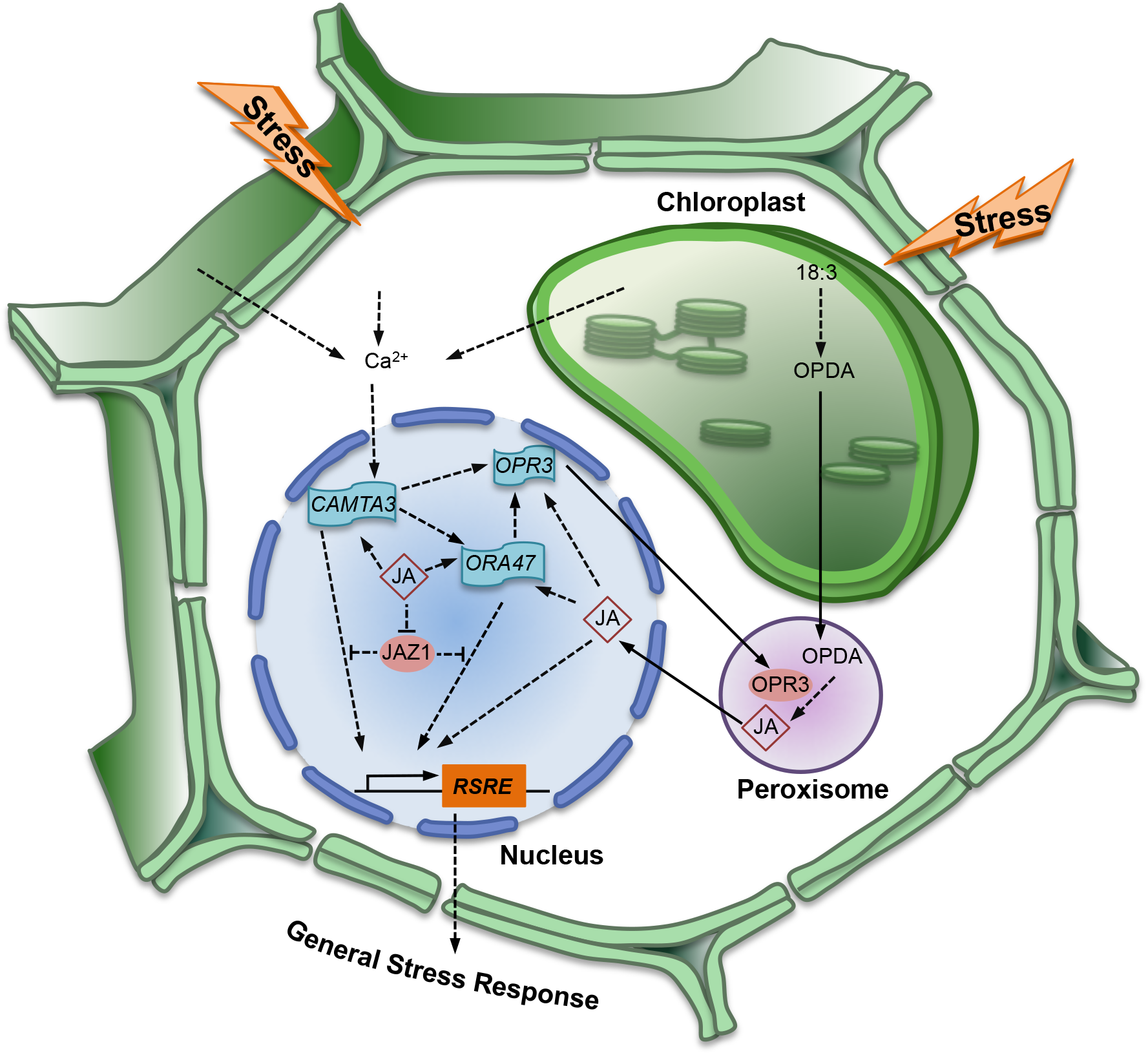
Schematic model depicting the intraorganellar cooperativity enabling the coordinated and complex interplay between transcriptional regulators and JA biosynthesis and signaling cascade involved in the induction of a general stress transcriptional hub and response to fluctuating environment.

## Materials and Methods

### Plant material and growth condition

*Arabdopsis thaliana* seedlings were grown in 16-h light/8-h dark cycles at ~22 ° C on half Murashige and Skoog medium. 11-days-old *TPT-ORA47* and *TPT-ORA47/camta3* seedling were treated with β-estradiol (10 μM, 72hr). 14-days-old WT and *camta3* seedling were treated with β-estradiol (10 μM, 0hr) as the control. Two-week-old seedlings were treated with BAPTA (10 mM, 24hr), MeJA (10 μm, 90min), or mechanical wounding of a single leaf per plant by forceps as previously described (Benn *et al*., 2014, Benn *et al*., 2016, Jiang *et al*., 2018). The egg cellspecific promoter-controlled CRISPR/Cas9 system with the Golden Gate cloning method was used for obtaining CRISPR-*ora47* mutants as previously described (Wang *et al*., 2015). Sequences of the two CRISPR guide RNAs are listed in Table S1.

### Yeast one-hybrid assay

Enhanced yeast one-hybrid (eY1H) assay was employed for screening of TFs binding to RSRE as previously described (Pruneda-Paz *et al*., 2009, Gaudinier *et al*., 2011). Specifically, the TaKaRa Gold Yeast one-hybrid library screening system (#630491) was used for screening RSRE motif binding proteins. The sequence of RSRE motif (the “DNA bait”) is cloned to create the 4xRSRE::LacZ and 4xRSRE::Luciferase (LUC) reporter constructs. The X-gal was used to detect the LacZ activity. The charge-coupled device (CCD) camera (Andor Technology DU-434BV) was used to detect Luciferase activity signals. Images were acquired every 5 min for 2h. The Andor Solis Software (version 14) was used to quantify the Luciferase activity as previously described (Pruneda-Paz *et al*., 2009, Pruneda-Paz and Kay, 2010, Benn *et al*., 2014, Pruneda-Paz *et al*., 2014, Breton *et al*., 2016).

### Luciferase-Activity quantification

The CCD camera (Andor Technology DU-434BV) was used to detect Luciferase activity signals. Specifically, plants were sprayed with 1.0 mM luciferin (Promega) in 0.1% Trion X-100. Images were acquired every 5 min for 4h for Arabidopsis, and every 15 min for 4h for tobacco. Quantification of *RSRE::LUC* activity were performed for a defined area as mean counts pixel^-1^ exposure time^-1^ by using the Andor Solis Software (version 14) as previously described (Benn *et al*., 2014). ~30-50 plants per genotypes per treatment are used as biological replicates.

### Plant hormone extraction and quantification

14-days-old plants grown in 16-h light/8-h dark cycles at ~22 ° C on half Murashige and Skoog medium were harvested and grinded in liquid nitrogen. Samples were then stored in −80 °C. Each genotype had four to five biological replicates. The weight of each replicate is 50 mg. Extraction and quantification of OPDA and JA by using the gas chromatography-mass spectrometry (GC-MS) were performed as previously described (Savchenko et al., 2010).

### Quantification of gene expression

Total RNA was isolated by using the Aurum total RNA mini kit (Bio-Rad) and treated with DNase (Bio-Rad) to avoid DNA contamination. One microgram of RNA was reverse transcribed by using iScript cDNA Synthesis Kit (Bio-Rad). 10 μl SsoAdvanced Universal SYBR Green Supermix reagents (Bio-Rad) per reaction (20 μl total volume), and the CFX96 real-time PCR detection system (Bio-Rad) were used to perform the Real time PCRs. (Walley *et al*., 2007). AT4G26410 (M3E9) was used as the control gene. Each experiment was performed with three biological and three technical replicates. Two-tailed Student’s *t* tests were performed for two-group samples, and ANOVA tests with Tukey’s honest significant difference (HSD) tests were performed for more than two-group samples. Table S1 listed all used primer sequences.

### Agro-infiltration-based transient assays in *Nicotiana benthamiana*

*N. benthamiana* transient assay was used to examine potential role of JAZ1 in regulation of ORA47- and CAMTA3-activated RSRE::LUC activity. Specifically, pENTR/D-TOPO (Invitrogen) and Gateway systems were used to assemble *35S::JAZ1, 35S::ORA47, 35S::CAMTA3* and RSRE::LUC constructs. The constructs were introduced into *Agrobacterium* GV3101 and subsequently used for infiltration of *N. benthamiana* leaves by using 1ml syringe, followed by luciferase activity signal detection using CCD camera after 48 hours of infiltration (Benn *et al*., 2014). 20 leaves are used as biological replicates for signaling calculation and analyses.

## Accession Numbers

*ORA47* (AT1G74930), *CAMTA3* (AT2G22300), *OPR3* (AT2G06050) and *JAZ1* (AT1G19180).

## Acknowledgements

This work was supported by Dr. John W. Leibacher and Mrs. Kathy Cookson endowed chair funds to KD, and by grant from the National Science foundation (NSF Award No: 1755452) to JPP, and National Institutes of Health (NIH; R01GM107311-8) and National Science foundation (NSF Award No: 2104365) to KD.

---

**Table S1.** Used primer sequences.

**Figure S1.**
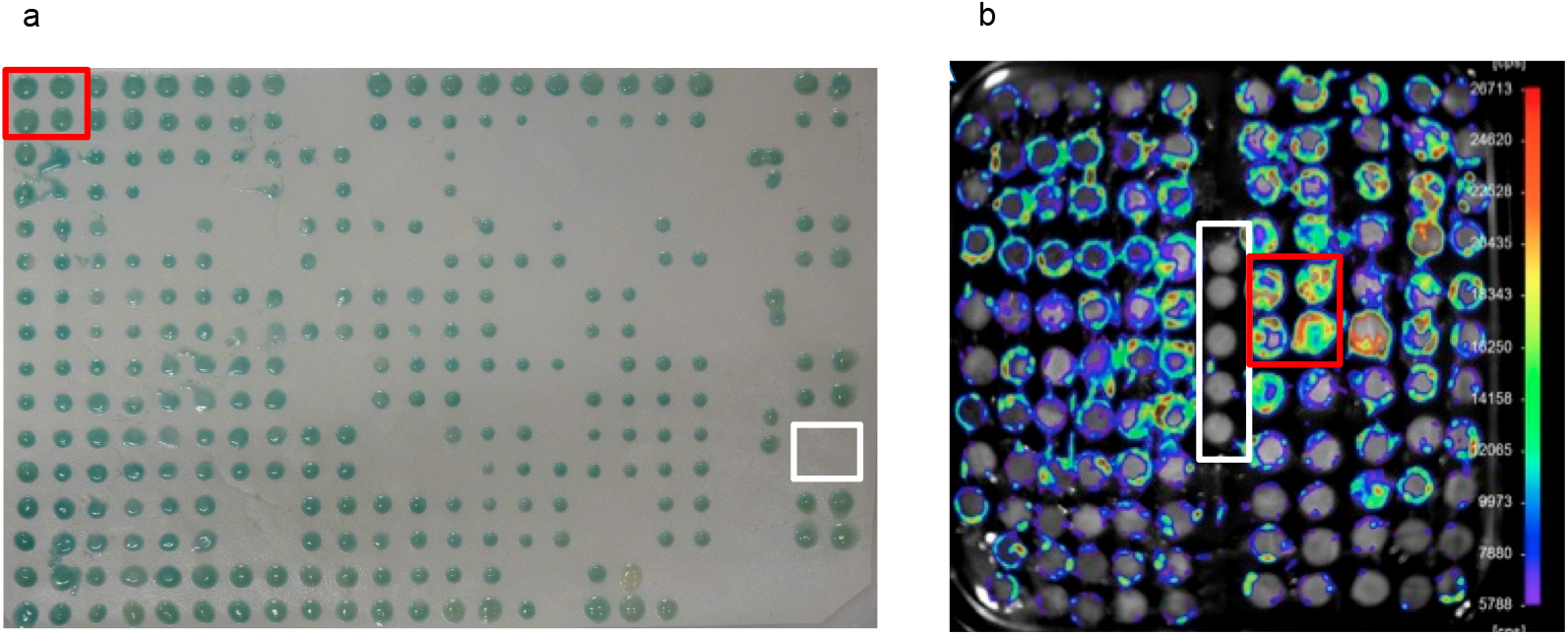
Identification of RSRE-binding proteins by Yeast-one-hybrid assay. Proteins binding to RSRE motif are identified by Y1H using lacZ (a) and luciferase (b) reporters. Each protein has four technical replicates, and the control has four technical replicates in Panel a, and five technical replicates in Panel b. ORA47 (red rectangle) and negative control (white) are marked.

**Figure S2.**
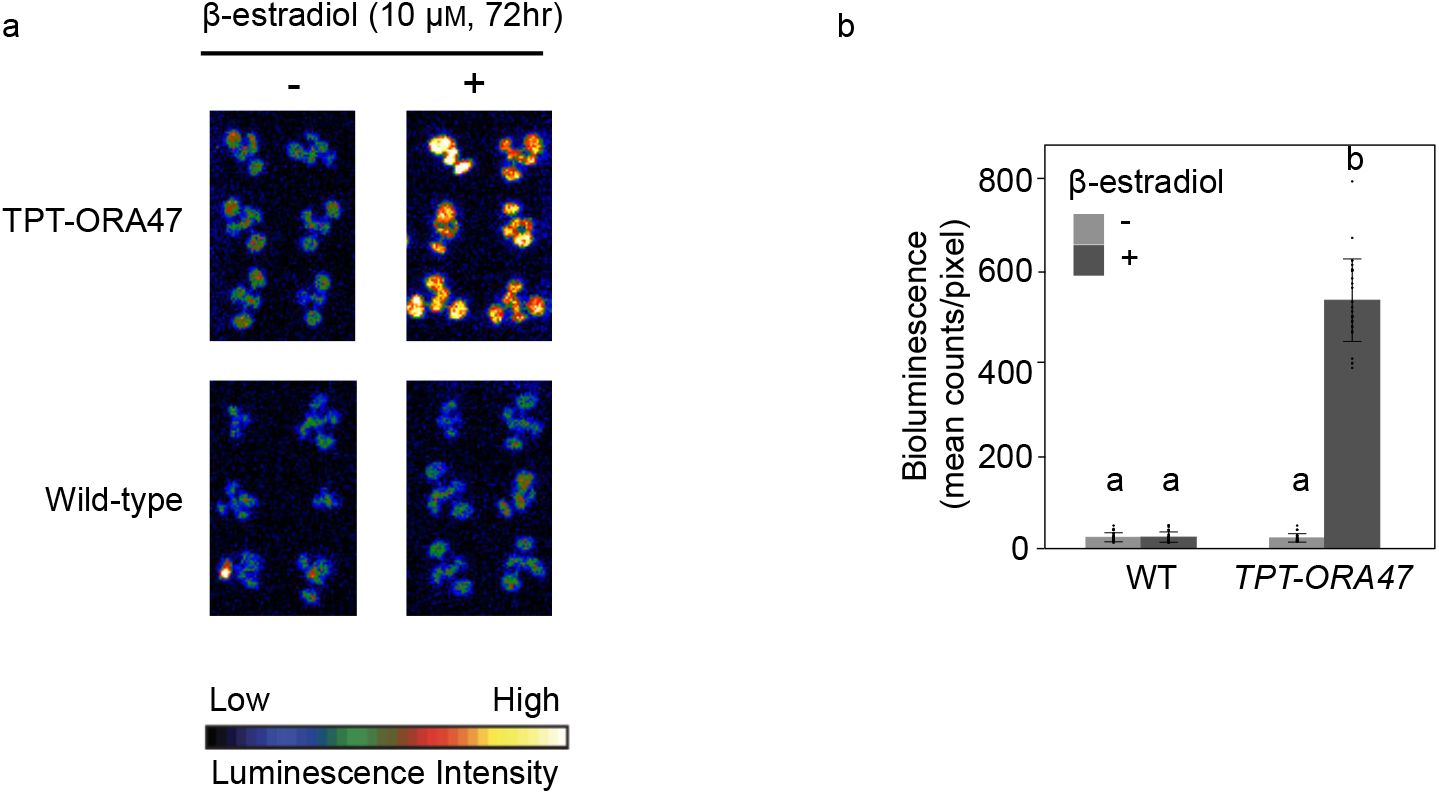
β-estradiol induces RSRE::LUC activity in TPT-ORA47 line. (a) Representative images of RSRE::LUC activity in control (-) and 72 hr post β-estradiol (10 μM) treatment (+) of wild type (WT) and inducible TPT-ORA47 lines. The color-coded bar displays the intensity of LUC activity. (b) Quantitative measurements of LUC activity of plants shown in Panel A. Bars that do not share a letter represent statistically significant differences (p<0.05) by ANOVA test with Tukey’s honest significant difference (HSD) test. 30 plants per genotype per treatment were used as biological replicates. The error bar is the standard deviation of biological replicates.

**Figure S3.**
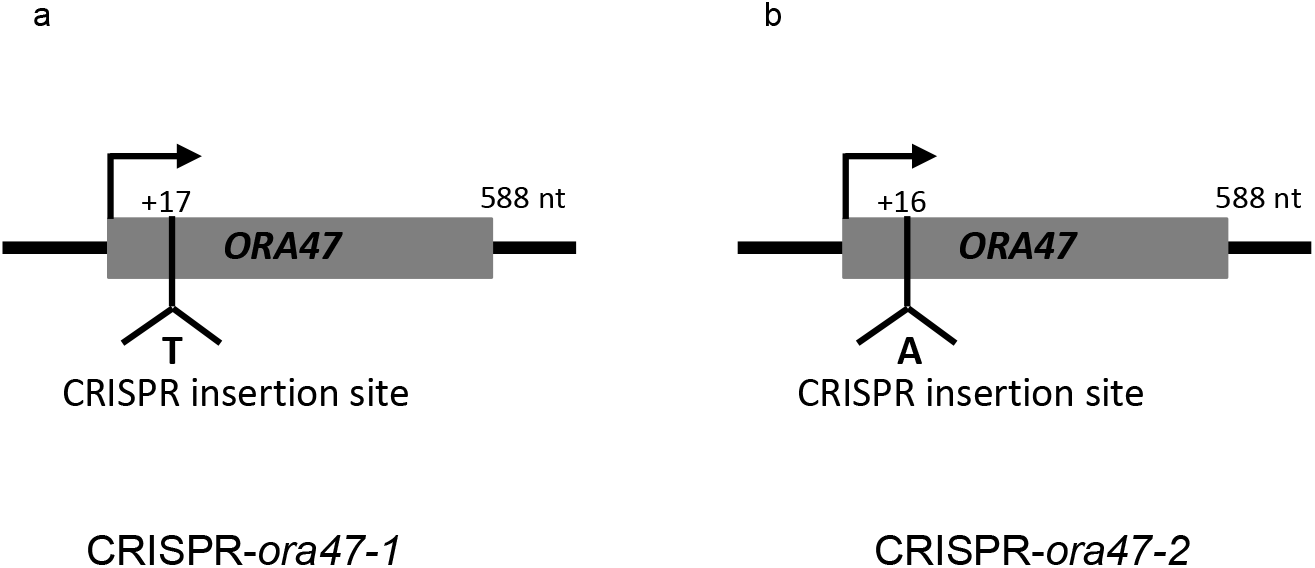
The schematic presentations of two independent CRISPR insertion sites in the *ORA47* resulting in generation of the two CRISPR-*ora47-1* and −*2* lines. Used primer sequences are shown in Table S1.

**Figure S4.**
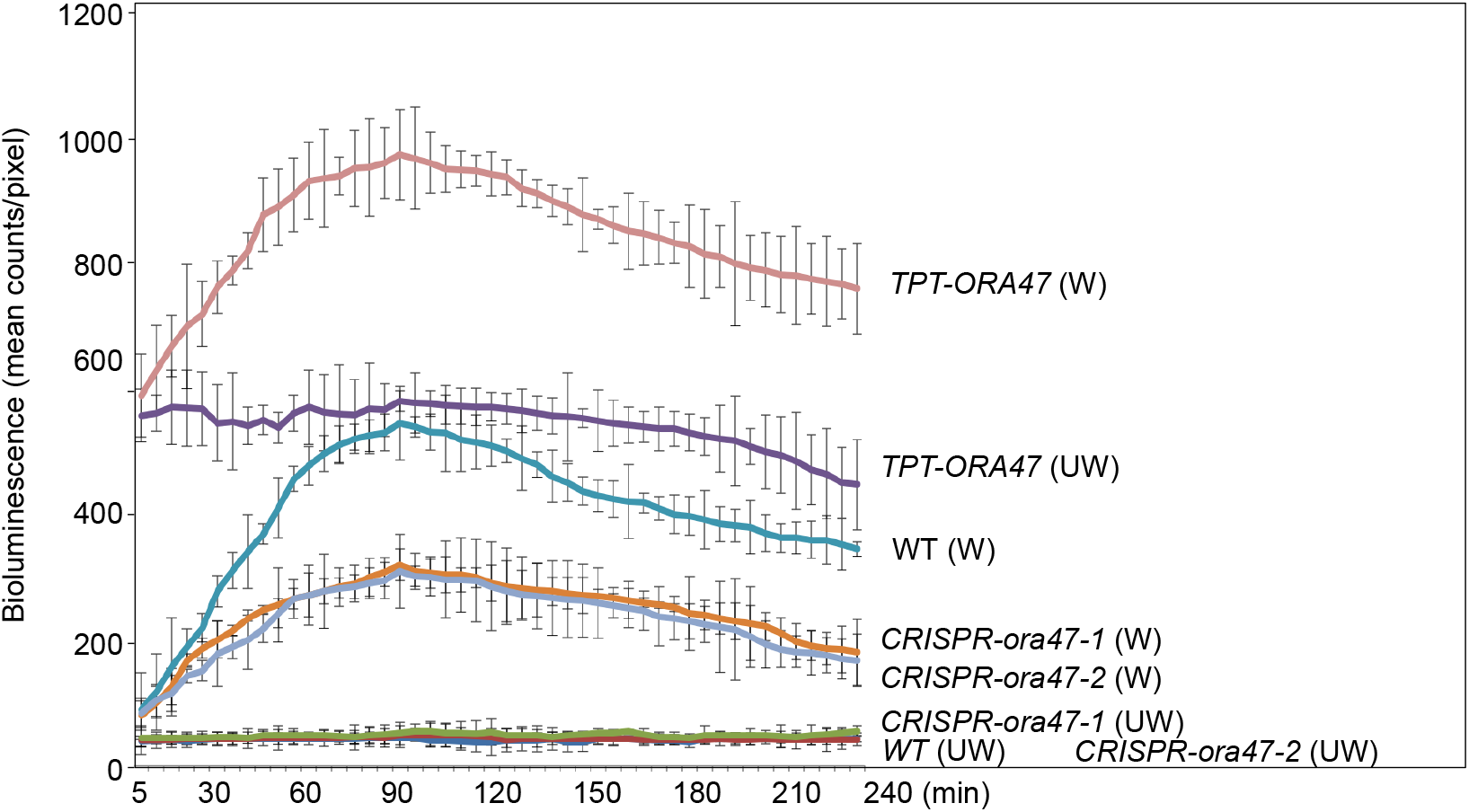
Time course of RSRE-driven LUC activity in control (UW) and wounded (W) genotypes. LUC activity detected 5 min post mechanical wounding, peaks at 90 min post wounding, as shown by signal intensity measurements every 5 min for 4 h in unwounded (UW) and wounded (W) WT, TPT-ORA47, CRISPRora47-1 and CRISPR-ora47-2 lines. The average signals obtained from 36 individual plants at each time point were used for the line plot. The error bar is the standard deviation of signals in biological replicates.

